# A habitat class to land cover translation model for mapping Area of Habitat of terrestrial vertebrates

**DOI:** 10.1101/2021.06.08.447053

**Authors:** Maria Lumbierres, Prabhat Raj Dahal, Moreno Di Marco, Stuart H. M. Butchart, Paul F. Donald, Carlo Rondinini

## Abstract

Area of Habitat (AOH) is defined as the ‘habitat available to a species, that is, habitat within its range’ and is produced by subtracting areas of unsuitable land cover and elevation from the range. Habitat associations are documented using the IUCN Habitats Classification Scheme, and unvalidated expert opinion has been used so far to match habitat to land-cover classes generating a source of uncertainty in AOH maps. We develop a data-driven method to translate IUCN habitat classes to land-cover based on point locality data for 6,986 species of terrestrial mammals, birds, amphibians and reptiles. We extracted the land-cover class at each point locality and matched it to the IUCN habitat class(es) assigned to each species occurring there. Then we modelled each land cover class as a function of IUCN habitat using logistic regression models. The resulting odds ratios were used to assess the strength of the association of each habitat land-cover class. We then compared the performance of our data-driven model with those from a published expert knowledge translation table. The results show that some habitats (e.g. forest and desert) could be more confidently assigned to land-cover classes than others (e.g. wetlands and artificial). We calculated the association between habitat classes and land-cover classes as a continuous variable, but to map AOH, which is in the form of a binary presence/absence, it is necessary to apply a threshold of association. This can be chosen by the user according to the required balance between omission and commission errors. We demonstrate that a data-driven translation model and expert knowledge perform equally well, but the model provides greater standardization, objectivity and repeatability. Furthermore, this approach allows greater flexibility in the use of the results and allows uncertainty to be quantified. Our model can be developed regionally or for different taxonomic groups.

## INTRODUCTION

Habitat loss is the most important driver of biodiversity decline (Díaz et al.,, 2019). Therefore, there is an urgent need to determine where habitat is located within each species’ distribution (Pimm et al., 2014; Brooks et al., 2019). Several approaches have been developed to map global species’ distributions, but accurate spatial data are only available for a limited number of species (Rondinini et al., 2005; Rondinini & Boitani 2012).

The most complete dataset of maps of species’ ranges is that available in the International Union for Conservation of Nature (IUCN) Red List (www.iucnredlist.org). The IUCN Red List has assessed more than 134,400 species, and species groups, including mammals, amphibians, and birds, have been comprehensively assessed. The IUCN range maps are generally drawn to minimize errors of omission (i.e. false absence), with the result that they often contain substantial areas that are not occupied by the species, and so suffer from errors of commission (i.e. false presence) (Ficetola et al., 2014; Di Marco et al., 2017).

Area of Habitat (AOH; previously known as extent of suitable habitat, or ESH) is the ‘habitat available to a species, that is, habitat within its range’ (Brooks et al., 2019). AOH maps are produced by subtracting unsuitable areas from range maps, using data on each species’ associations with land cover and altitude (Beresford et al., 2011; Rondinini et al., 2011; Ficetola et al., 2015), and attempts to reduce commission errors in range maps. Therefore, the production of AOH maps requires an understanding of which habitats a species occurs in and where those habitats are located within its range.

Information on habitat preferences is documented for each species assessed on the IUCN Red List (IUCN 2013) following the IUCN Habitats Classification Scheme (IUCN habitat; IUCN, 2012), a classification and coding system of habitats that ensures global consistency. IUCN standardized habitat definitions independently of taxonomy or geography. However, IUCN habitat classes are not spatially explicit, although recent efforts have attempted to delimit them (Jung et al., 2020). Land-cover classes derived from remote sensing have been widely used as a surrogate of habitat (e.g. Buchanan et al., 2008; Beresford et al., 2011; Rondinini et al., 2011; Tomaselli et al., 2013; Montesino Pouzols et al., 2014; Corbane et al., 2015; Santini et al., 2019), although habitat is a complex multi-dimensional concept that is difficult to simplify into land-cover classes.

A translation table between habitat and land-cover classes is typically used to represent IUCN habitat classes spatially and to produce AOH maps. This is a table that shows which habitat classes map onto which land-cover classes. Previous versions have been based solely on expert knowledge, raising concerns about the accuracy and objectivity of the resulting associations, as the assumptions generated in the translation process are rarely considered in detail and the errors are difficult or impossible to quantify (Bradley et al., 2012). Furthermore, there is a lack of standardization in the procedure (Seoane et al., 2005), which is subject to variability in expert opinion (Johnson & Gillingham, 2004).

Repositories of point locality data (i.e. locational records where particular species have been recorded (Rondinini et al., 2006)) primarily from citizen science have been successfully used in habitat suitability models (e.g. Gueta & Carmel, 2016; Bradter et al.,, 2018; Crawford, Olds, Maerz, & Moore, 2020). The potential, therefore, exists to use such data also to develop an objective data-driven translation table between habitat and land-cover classes by extracting information on land cover from point localities of species with different habitat associations.

Here we propose a standardized, data-driven methodology to produce a translation table between IUCN habitat classes and two widely used global land-cover maps, the Copernicus Global Land Service Land Cover (CGLS-LC100; Copernicus Global Land Operations “Vegetation and Energy”, 2018a; Buchhorn et al., 2019) and the European Space Agency Climate Change Initiative land cover 2015 (ESA-CCI; ESA, 2017) using point locality data for mammals, birds, amphibians and reptiles (the best-documented groups of species). The aim of this analysis was to develop a translation table that quantifies the power of association between land cover and habitat classes. In doing so, we aim to illustrate a method that improves on expert opinion by (a) quantifying errors in associations between habitat and land cover classes, (b) being flexible to the needs of the user in terms of the required trade-off between reducing commission errors and increasing omission errors, and (c) can be developed at different spatial scales, for different taxa, using any set of habitat or land-cover classes.

## METHODS

### Data cleaning and preparation

We downloaded point locality data for mammals (GBIF, 2019; GBIF, 2020), amphibians (GBIF, 2020) and reptiles (GBIF, 2020) from the Global Biodiversity Information Facility (GBIF) and for birds from GBIF (GBIF, 2019; GBIF, 2020) and eBird (eBird Basic Dataset, 2019). The data were restricted to point localities dated from January 2005 to December 2018 for the model building (70% training and 30% test), and from January 2019 to December 2020 for the evaluation of the model. For eBird data, we selected only stationary point localities with a coordinate uncertainty of less than 30 m. To minimize errors, and uncertainties inherent to repositories of point locality data, we included only the most precisely georeferenced points (Rondinini et al., 2006; Meyer 2012) and applied a set of filters following the guidelines of Boitani et al., (2011). The main attributes considered were currency, spatial accuracy and spatial coverage (Fig. 1).

**Figure 1.**
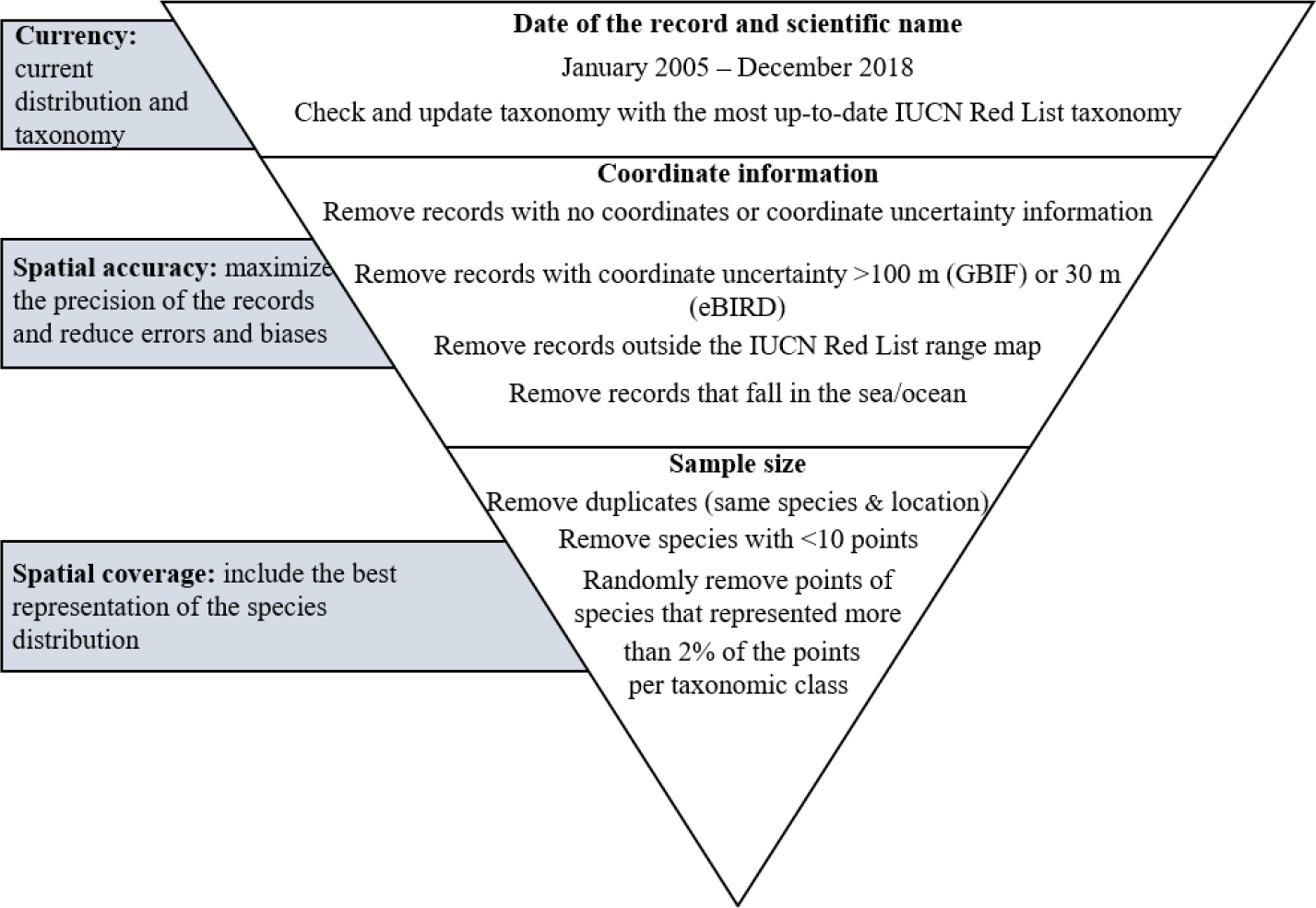
Description of the point locality cleaning process, following Boitani et al., (2011). The factors considered were currency, spatial accuracy and spatial coverage. The filters were applied from top to bottom.

To make it clear where we are referring to explicit classes, we present land-cover class names in quotation marks and IUCN habitat class names in italics.

The habitat class(es) association of each species were extracted from the IUCN habitat (IUCN, 2020). These follow a hierarchical classification of habitat with three levels. The definitions consider biogeography, latitudinal zonation and depth in marine systems. In this analysis, we used Level-1 habitat classes for all habitats except for artificial terrestrial, for which we used a modification of Level-2 (Appendix S1). We subdivided artificial terrestrial into three subclasses because in terms of land cover these are distinct habitat classes that could aggregate different species (Ducatez et al., 2018).

Because the land-cover classes from the two remote sensing products are exclusively terrestrial, we limited the analysis to species coded only to terrestrial habitat classes, thus excluding species coded to one or more IUCN marine habitats. We also excluded species coded to more than five Level-1 habitat classes, because habitat generalists are likely to add little information to the habitat-land cover relationship. In contrast, specialist species coded to only one habitat class provide more insight into the relationship between habitat and land cover. For that reason, for each taxonomic class, we randomly subsampled point records from species coded to more than one habitat class to match the number of points of species coded to one habitat and thereby gave equal weight to habitat specialists even when they had fewer points.

We developed models for two different global land-cover products derived from remote sensing: CGLS-LC100 and ESA-CCI. The CGLS-LC100 has a 100-m spatial resolution and a global classification accuracy of 80.2% (Copernicus Global Land Operations “Vegetation and Energy”, 2018b). The ESA-CCI has a 300-m spatial resolution and a global classification accuracy of 71.1% (ESA 2017). It is part of a time series from 1992 to 2015, of which we used the 2015 map. Both products use the United Nations Food and Agriculture Organization Land Cover Classification System (UN-LCCS), although they have different legends. CGLS-LC100 has 12 land-cover classes at Level-1 and 23 classes at Level-3 (Level-2 is not used by CLGS-LC100), and we used Level-3. ESA-CCI has 22 land-cover classes at Level-1 and 38 classes at Level-2. In this analysis, we only used Level-1 because Level-2 is only available for some regions of the globe.

To prepare the data for the model, we extracted the land-cover class at the coordinates of each point locality. Some land-cover classes did not have enough point localities falling within them to be modeled, although in all cases these were land-cover classes with very low global coverage. For CGLS-LC100, the under-represented land-cover classes were “open forest deciduous needle leaf” (10 points, 0.03% of global land surface), “snow and ice” (108 points, 3.1% of global land surface), “moss and lichen” (124 points, 2.3% of global land surface) and “closed forest deciduous needle leaf” (383 points, 3.0% of global land surface). For ESA-CCI, the only class represented too infrequently for analysis was “lichens and mosses” (713 points, 2.2% of global land surface).

### Modeling of habitat-land cover associations

To quantify the relationship between IUCN habitat classes and land-cover classes, we modeled the presence or absence of each land-cover class as a function of the IUCN habitat class(es) of the species whose point localities fell within it. An important consideration for modeling was that the number of habitat classes per species varied from one to five. Therefore, it was impossible to model land-cover class as a one-to-one relationship with habitat classes, as each point location was associated with one or multiple habitats. This consideration restricted the number of models we could use for our analysis. We required a flexible model that allowed a many-to-many match between habitat classes and land-cover classes to model this matrix of habitat vs land-cover class relations. In multinomial logistic regression models, the data and the computational power requirements increase exponentially with the number of response categories. In our case, with more than 20 land-cover categories, this option was not feasible. Therefore, we modeled each land-cover class separately, transforming it into a binary variable of 1 or 0 (land cover present/not present in a point locality). Then, we used logistic regressions to model the binary land-cover class variable as a function of the different habitat classes:

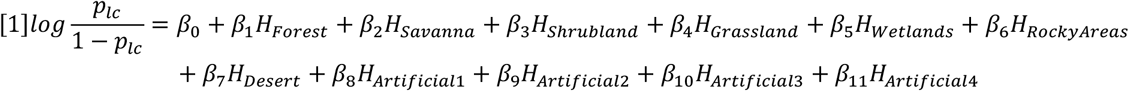

where (p_lc_/(1-p_lc_)) is the land cover odds ratio and β_x_ are the model parameters for each of the habitats *H*_x_.

The transformation of the land-cover class into a binary form for each of the models generated a highly unbalanced variable, with many more zeroes than ones. In a logistic regression model, unbalanced data underestimate the probability of an event so it is recommended to adjust the number of 1s and 0s (King & Zeng 2001; Pozzolo et al., 2015). We therefore randomly subsampled the 0s in the training set before running the model. The assumption behind this is that in the majority class there are many redundant observations and randomly removing some of them does not change the estimation of the within-class distribution (Pozzolo et al., 2015).

To reduce the intrinsic spatial and taxonomic bias point locality data (Boitani et al., 2011; Meyer et al., 2016), and to account for multiple but varying numbers of point localities per species, we added taxonomic and spatial variables as random effects in the model (Bird et al., 2014). As taxonomic variables, we added species nested within taxonomic class (Amphibia, Reptilia, Aves, Mammalia). Adding intermediate taxonomic groupings (e.g. family or genus) in the nesting would result in many factor levels with single or very few replicates. To test whether there was any bias between taxonomic classes, we first produced separate models for each class, and found that the association between land cover and habitat classes from the different translation tables were very similar; therefore, we decided to model all classes together. As a spatial variable, we added the country of the point record as a random effect.

We used the coefficients of the models to evaluate the association between land-cover class and habitat classes. The intercept did not provide any information on the relationship between land-cover class and habitat class as it represents the odds of a point locality falling within a particular land-cover class after the subsampling of the data set, independently of the habitat (Ranganathan et al., 2017). The coefficients represent the odds ratio, in other words, the odds of a point locality falling in a particular land-cover class (when the species to which the point locality relates is coded for a particular habitat class) divided by the odds of the species occurring in that land-cover class when it is not coded for that habitat class. The ratio, therefore, indicates the extent to which being coded to a particular habitat class increases or decreases the odds of a species being found in a particular land-cover class. The units of the logit function are log(odds ratio), but for easier interpretation, we exponentiated them and present the results as odds ratios.

Odds ratio values below 1 indicate a negative association between land cover and habitat classes, while those above 1 indicate a positive association. As the odds ratio is a continuous variable, it is necessary to set a threshold to transform the results into a binary translation table that can be used to assign, or not, a particular habitat class to a particular land-cover class. The threshold can be modified according to the needs of the user based on the required balance between minimizing commission errors (land-cover classes incorrectly associated with a habitat class) and increasing omission errors (land-cover classes incorrectly omitted from a habitat class). Coefficients that had *p*-values higher than 0.05 were considered to indicate a lack of association between land cover and habitat classes. To adjust the significance threshold of the *p*-values for multivariable analysis, we used Bonferroni corrections.

To validate the models, we set aside 30% of the point occurrence data for testing, leaving 70% to train the model. As a validation test, we used the Area Under the Curve (AUC) from a Receiver Operating Characteristic (ROC) curve. AUC is a model accuracy measure that provides information on how well a model can distinguish among classes. In our case, we used it to test how well the models predicted the presence/absence of a point locally in a given land cover class. AUC values range from 0 to 1, a value of 0.5 means that the model does not performs better than random, while a value of 1 indicates that the model can perfectly separate the two groups.

We then compared the performance of the data-driven translation table with that of an expert knowledge translation table (Santini et al., 2019) based on the same ESA-CCI land cover classification used here. We did not find any published translation table that used CGLS-LC100. Santini et al., matched the ESA CCI land cover classes against Level-2 IUCN habitat classes, so we aggregated the habitat classes to Level-1 IUCN habitat classes to make the two translation tables comparable. We limited the comparison to birds and mammals because they were the taxonomic groups considered by Santini et al., For each species we mapped suitable habitat based on both tables. We assessed the proportion of points located in the suitable habitat (point prevalence) and compared it with the proportion of suitable habitat inside the species’ range (model prevalence) to determine whether the results were better than a randomly assigned set of points (Rondinini et al., 2011).

## RESULTS

The number of point localities and species available for this analysis was 200,683 and 455 respectively for mammals, 4,083,510 and 5,154 for birds, 92,327 and 479 for amphibians, and 131,077 and 898 for reptiles. For the CGLS-LC100 land-cover product, 71 coefficients showed a significantly positive association (odds ratio >1) and 38 coefficients showed a significantly negative association (odds ratio <1) between land-cover classes and habitat classes (Fig. 2). For the ESA-CCI land-cover product, 101 coefficients showed a significantly positive association and 40 coefficients showed a significantly negative association (Fig. 3).

**Figure 2.**
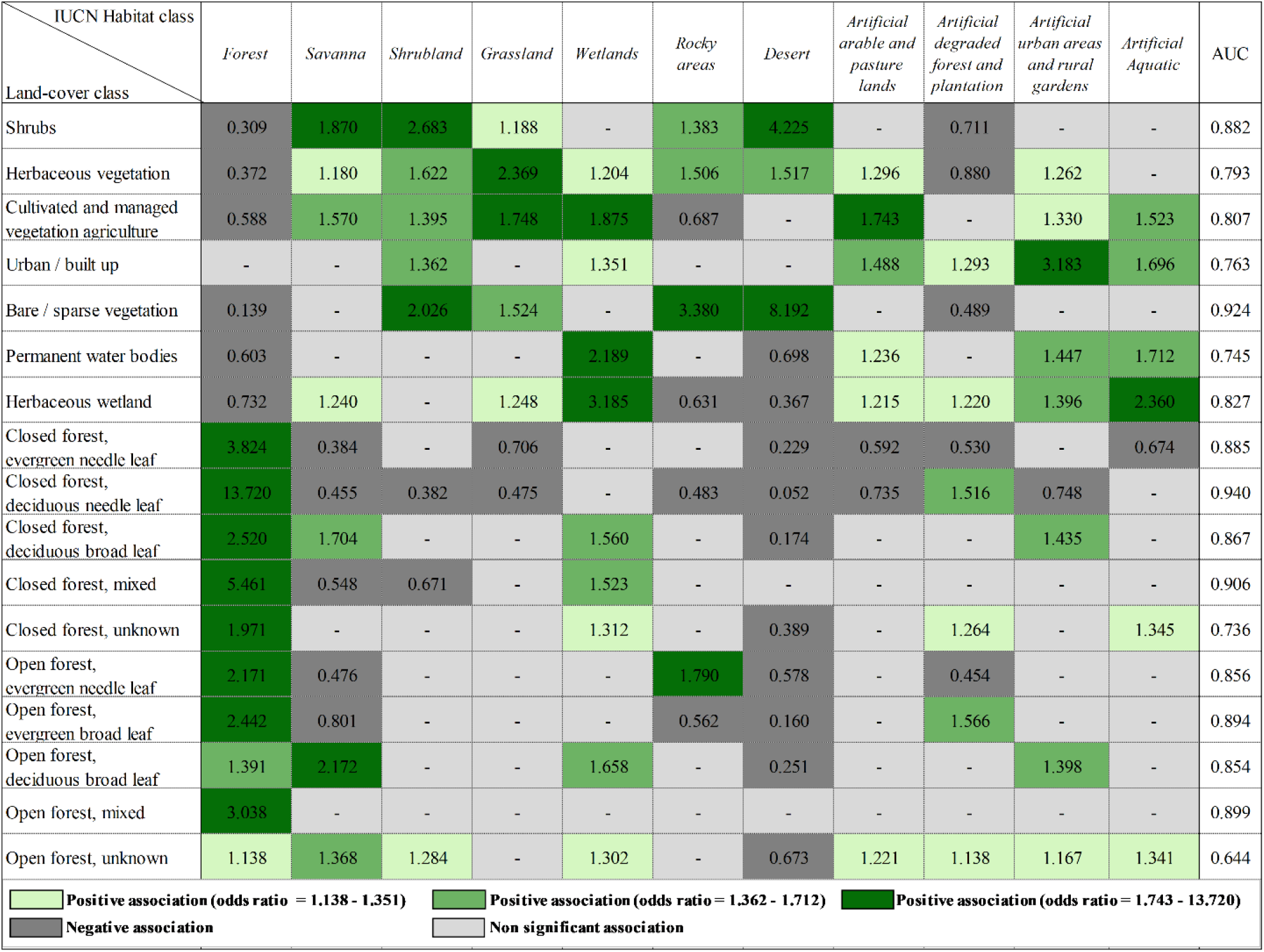
Odds ratio values describing the association between CGLS-LC100 land-cover classes and IUCN habitat classes. Odds ratio values significantly < 1 indicate a negative association, and values significantly > 1 indicate a positive association. The significantly positive associations are divided into tertiles (shown in shades of green), indicating three possible options for setting a threshold to convert continuous variables into a binary association/non-association variable for creating AOH maps. AUC indicates the values of Area Under the Curve from a ROC, a measure of accuracy of a classification model.

**Figure 3.**
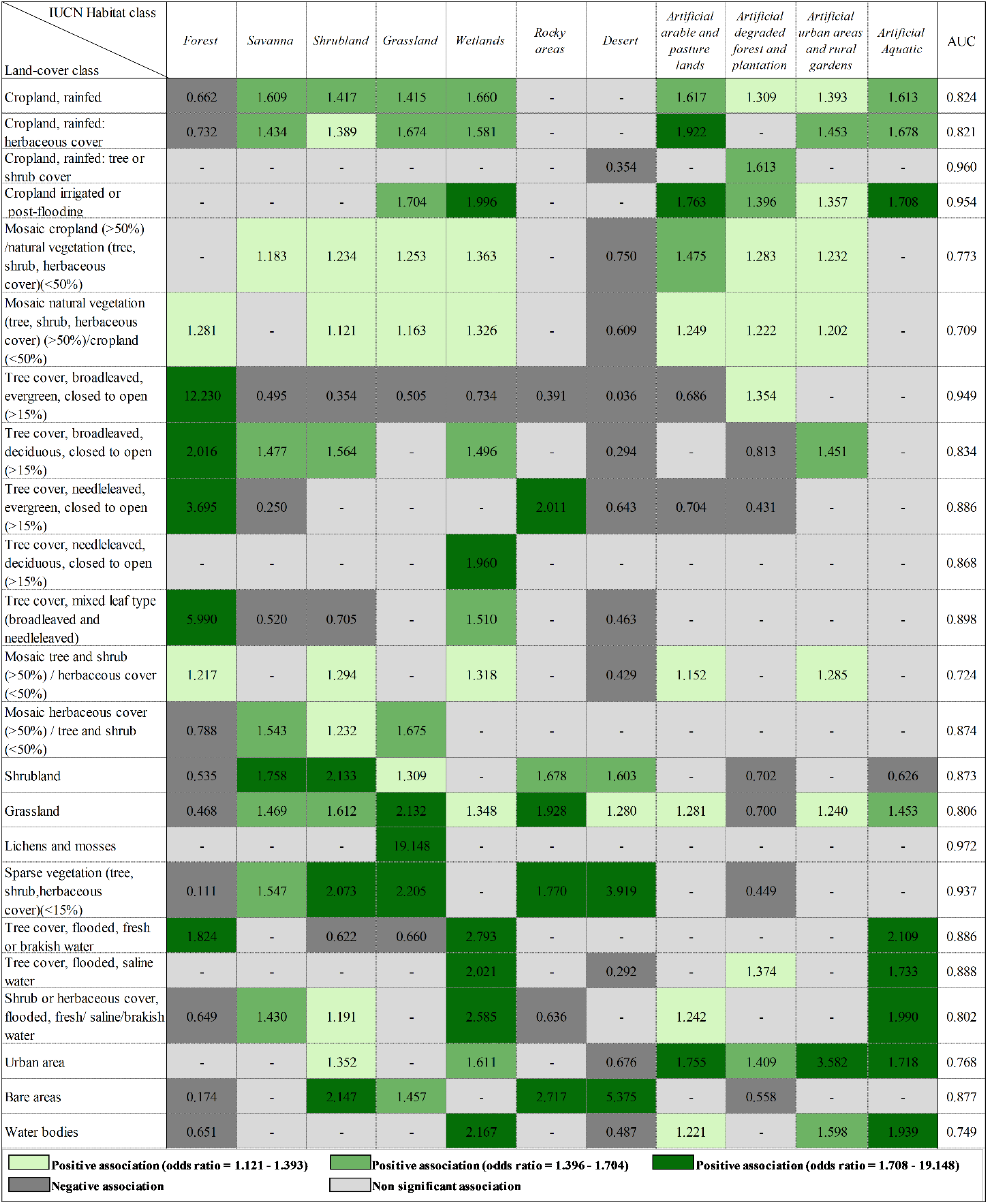
Odds ratio values describing the association between ESA-CCI land-cover classes and IUCN habitat classes. Odds ratio values significantly < 1 indicate a negative association, values significantly > 1 indicate a positive association. The positive associations are divided into tertiles (shown in shades of green), indicating three possible options for setting a threshold to convert continuous variables into a binary association/non-association variable for creating AOH maps. AUC indicates the values of Area Under the Curve from a ROC, a measure of accuracy of a classification model.

Higher odds ratios (>1) indicated stronger positive associations between land cover and habitat classes, and lower odds ratios (nearer to zero) indicated stronger negative associations. We divided the significantly positive values into tertiles to identify three potential thresholds for creating a table of binary association/non-association variables for producing AOH maps: 1.138-1.351, 1.362-1.712 and 1.743-13.720 for CGLS-LC100, and 1.121-1.393, 1.396-1.704 and 1.708-19.148 for ESA-CCI.

*Forest* and *Desert* had the strongest positive associations between land cover and habitat classes. The *Forest* habitat class was associated with almost all the forest and tree cover land-cover classes (CGLS-LC100 average positive odds ratio = 3.8; ESA-CCI average positive odds ratio = 4.0) and with no other land-cover classes. The *desert* habitat class was also strongly associated with particular land-cover classes: “shrubs”, “herbaceous vegetation”, and “bare/sparse vegetation” in CGLS-LC100 (average positive odds ratio = 4.6) and “shrubland”, “grassland”, “sparse vegetation (tree, shrub, herbaceous cover < 15%)” and “bare areas” in ESA-CCI (average positive odds ratio = 3.0). *Rocky areas* were associated with almost the same land-cover classes as *Desert* but with lower odds ratios.

*Savanna*, *Shrubland* and G*rassland* habitat classes were associated with “shrubs”, “herbaceous vegetation” and “cultivated and managed vegetation agriculture” in CGLS-LC100 land cover, and “cropland”, “herbaceous cover”, “shrubland”, “grassland”, “sparse vegetation”, “mosaic cropland” and “mosaic herbaceous cover” in ESA-CCI. However, the power of association varied between these different combinations. The *Savanna* habitat class was also associated with some forest classes while *Shrubland* and *Grassland* habitats were also associated with bare areas.

We divided artificial terrestrial habitats into three different classes: *Artificial arable and pasture lands*, *Artificial degraded forest and plantations*, and *Artificial urban and rural gardens*. These habitats had the least certain relationships because the odds ratio values were the closest to 1 (CGLS-LC100 average positive odds ratio = 1.367, 1.333 and 1.577 respectively; ESA-CCI average positive odds ratio = 1.468, 1.370 and 1.579 respectively). Some unexpected land-cover classes were associated with these habitat classes, e.g. *Arable and pasture lands* and *degraded forest and plantations* were associated with “urban areas”. However, these unexpected associations disappeared when increasing the threshold.

*Wetland* and *Artificial aquatic* habitats had intermediate odds ratio values (CGLS-LC100 average positive odds ratio = 1.7; ESA-CCI average positive odds ratio = 1.8). In terms of land-cover associations, they were associated (in some cases strongly) with land-cover classes related to water, but also to some land-cover classes that have no relation with wetlands or aquatic environments (e.g. some type of forest or cultivated areas).

The AUC of models for CGLS-LC100 ranged from 0.644 to 0.940. The land-cover classes with the lowest AUC were the “open and closed unknown forest” (AUC = 0.644 and 0.736) classes, followed by “water bodies” (AUC = 0.745) and “urban areas” (AUC = 0.763). Those with the highest AUC values were the other forest classes (AUC range 0.854 – 0.940) and “bare and sparse vegetation” (AUC = 0.924). For ESA-CCI, the AUC ranged from 0.709 to 0.972. The land-cover classes with the lowest AUC were mosaic land-cover classes (AUC range 0.709 - 0.874), followed by “water bodies” (AUC = 0.750) and “urban areas” (AUC = 0.768). The land-covers with the highest AUC values were “lichens and mosses” (AUC = 0.972), “cropland irrigated or post-flooding” (AUC = 0.954), “sparse vegetation” (AUC = 0.937) and tree cover land classes (AUC range 0.834 – 0.949).

The results of the models can also be mapped spatially (Fig. 4) using one of the three thresholds of associations between habitat and land-cover classes. In such maps, habitats are overlaid because the same land-cover class may represent more than one habitat class and/or because both habitats occur in the same geographical areas. The overlap among habitats increases as the threshold of association is reduced.

**Figure 4.**
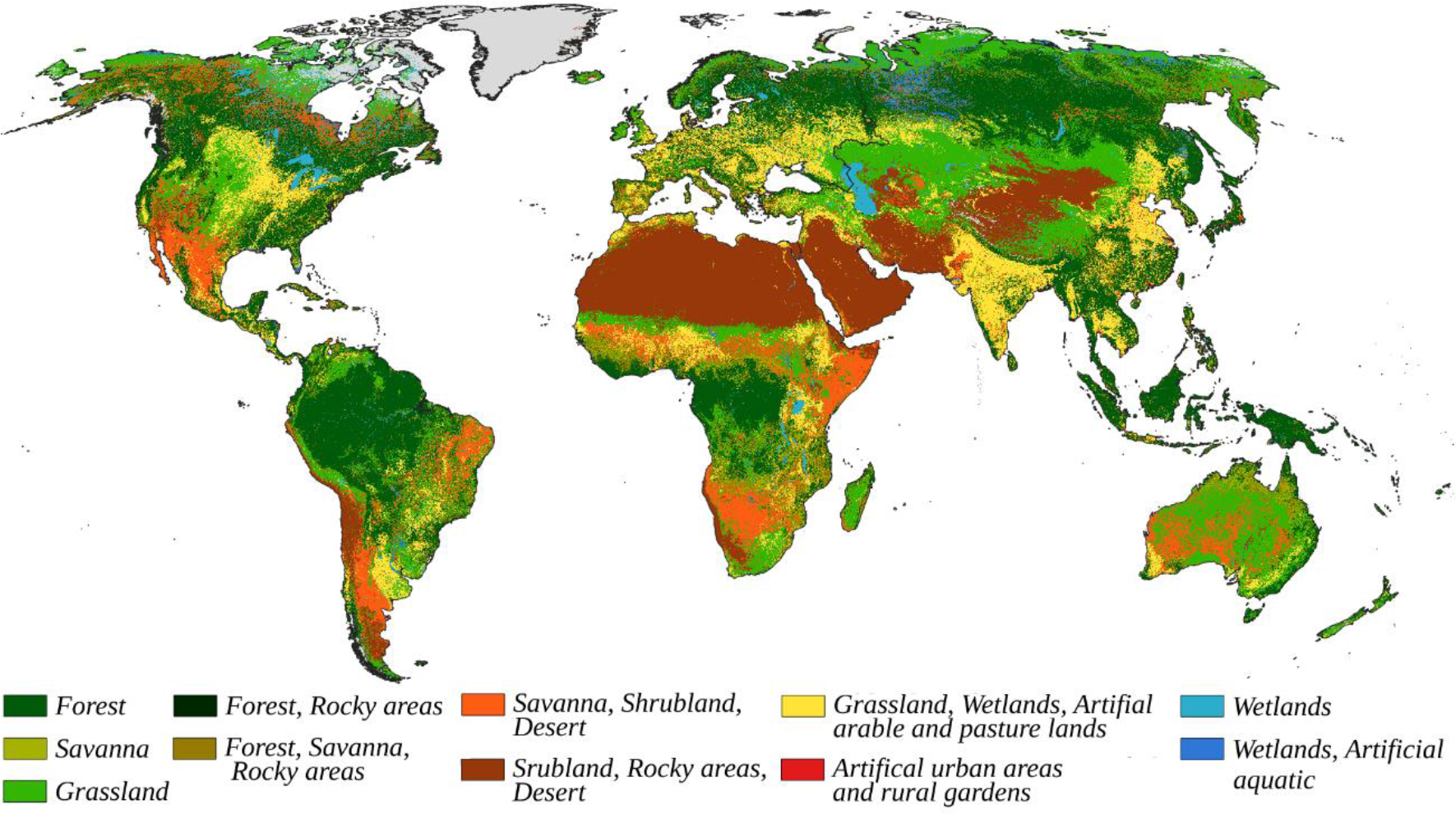
Map of habitat classes (Level 1) from the IUCN Habitat classification scheme based on the highest threshold for CGLS-LC100 data-derived translation (Fig 2).

To compare the performance of the data-driven table and the expert-derived table, we used 211,304 point localities for 489 species of mammal and 461,277 point localities for 2,112 species of bird. We compared the point prevalence (the proportion of georeferenced points falling in the land-cover classes assigned by each translation table according to the habitat of each species) between our data-driven method and the expert based assessment of Santini et al., and found that point prevalence in Santini et al., (2019) was similar to the point prevalence we found from our model when using the middle and high odd-ratio thresholds (Table 1). The ratio between point prevalence and model prevalence (the proportion of the range remaining after apparently unsuitable land cover classes are excluded) between the two methods was also very similar, and higher than 1, indicating that the habitat associations were better than random for both approaches.

**Table 1.** Mean point prevalence and model prevalence for birds and mammals using the three tertile thresholds for ESA CCI land cover derived from data-driven assessment (see Figure 4) and the expert knowledge-based assessment of Santini et al., (2019).

|  | Lower tertile threshold | Middle tertile threshold | Upper tertile threshold | Santini et al., (2019) |
| --- | --- | --- | --- | --- |
| BIRDS |  |  |  |  |
| Point prevalence | 0.94 | 0.81 | 0.66 | 0.74 |
| Model prevalence | 0.91 | 0.76 | 0.59 | 0.68 |
| MAMMALS |  |  |  |  |
| Point prevalence | 0.93 | 0.82 | 0.67 | 0.73 |
| Model prevalence | 0.90 | 0.80 | 0.62 | 0.70 |

## DISCUSSION

By modeling the relationship between IUCN habitat classes and the CGLS-LC100 and ESA-CCI land-cover classes, we generated two translation tables, quantifying the strength of association between habitat and land cover classes. The strength of association is represented by the odd ratio values, which indicate the extent to which a species being coded to a particular habitat class increases or decreases the odds of that species being found in a particular land-cover class. The relationship between IUCN habitat classes and land cover classes is expressed as a continuous variable.

Among habitat classes, *Forest*, *Desert* and *Rocky areas* have the strongest associations with land-cover classes, perhaps owing to the higher accuracy of the relevant land-cover classes. For both CGLS-LC100 and ESA-CCI, the highest classification accuracy classes are “forest”, “tree cover areas” and “bare soil”. Using a different approach based on a decision tree, Jung et al., (2020) found that *Forest* has the highest validation accuracy, although they obtained lower validation accuracy for *Rocky areas* and *Desert* habitat classes.

On the other hand, *Wetlands* and *Artificial* habitats are more difficult to represent using land-cover maps. Wetland-related land-cover classes have the lowest classification accuracy in both land-cover maps. From a remote sensing perspective, wetlands are difficult to map because they are highly dynamic, with rapid phenological changes through the year (Gallant, 2015; Lumbierres et al., 2017). Remote sensing products at a global scale cannot distinguish small ponds or temporary water bodies (Pekel et al., 2016; Klein et al., 2017). Therefore, wetland land-cover classes have more omission errors, and this has a direct impact on the results of our model.

Artificial land-cover classes are also difficult to map, as they tend to be more heterogeneous (Álvarez-Martínez et al., 2018), producing misclassifications among land-cover classes. Land-cover maps have difficulty separating artificial land-cover classes from natural ecosystems, e.g., plantation from forest, grassland from cropland, or lake from reservoir (Álvarez-Martínez et al., 2018). Overall, species richness and average abundance are often lower in artificial environments than in their natural equivalent, even if there is variation across different biogeographical contexts (Barlow et al., 2007; Newbold et al., 2015) and this introduces commission errors. Moreover, we found that artificial land-cover are associated with some natural habitat classes. This is likely a consequence of greater accessibility of these habitats, and hence disproportionate prevalence in citizen science data (Meyer et al., 2015). Because a high proportion of citizen science point location data are recorded in artificial land-cover classes, there is an increased probability that species primarily associated with natural habitats are reported there, so a data-driven method may associate some natural habitats with artificial land-cover classes.

There are several differences between the two land-cover layers used to produce the translation tables that could determine the use of the table. CGLS-LC100 has a resolution of 100 m while ESA-CCI has a coarser resolution of 300 m, also CGLS-LC100 has an overall classification accuracy of 80.2% compared with 71.1% for ESA-CCI. Moreover, CGLS-LC100 avoids using mosaics classes and in general, mapping less complex habitats is easier than more heterogeneous habitats (Corbane et al., 2015; Álvarez-Martínez et al., 2018). However, ESA-CCI has the advantage of being available as a longer time series, 1992-2020 for ESA-CCI vs 2015-2019 for CGLS-LC100, which allows studying habitat changes. For both land cover maps we excluded some land-cover classes because of the lack of point localities; we recommend adding these land cover classes manually when using the translation tables, according to the user’s needs.

The coding of habitats to each species on the IUCN Red List could introduce some noise to the modeling process. Coding is based on qualitative assessment by experts, and is therefore susceptible to individual biases (Brooks et al., 2019; Santini et al., 2019). The current version of the Habitat Classification Scheme on the IUCN website is described as a draft version. We, therefore, recommend that IUCN updates and improves this document and anticipate this would influence our odds ratio estimates.

Both the data-driven table and the expert knowledge translation table represented land cover distribution inside the range better than random. However, our data-driven approach presents several advantages compared to an expert knowledge approach. We present the relationship between IUCN habitat and land cover classes as a continuous variable, allowing greater flexibility in using the results. To use the model to produce AOH maps, the user is able to decide a threshold of association to transform the results into a binary table according to the required balance between omission and commission errors. Moreover, a data-driven approach allows us to quantify the uncertainty associated with the habitat to land cover association and could help to evaluate potential uncertainties in the AOH maps. This approach can be used to develop a translation table between any set of habitat codes and any set of land cover variables at a global or regional scale. As better data (species point and land-cover maps) become available, the translation table can be improved, assuring objectivity, standardization, and repeatability.

## ACKNOWLEDGEMENTS

This research is part of the Inspire4Nature Innovative Training Network, funded by the European Union’s Horizon 2020 research and innovation program under the Marie Skłodowska‐Curie grant agreement no. 766417. MDM acknowledges support from the MIUR Rita Levi MontalcinLevii programme.

## AUTHOR CONTRIBUTIONS

ML, CR, PFD, and SHMB conceived the study. ML and PFD develop the analysis. ML led the writing of the manuscript. All authors contributed to drafts and gave final approval for publication.

## Notes

### Competing Interest Statement

The authors have declared no competing interest.

